# Venice: A New Algorithm for Finding Marker Genes in Single-Cell Transcriptomic Data

**DOI:** 10.1101/2020.11.16.384479

**Authors:** Hy Vuong, Thao Truong, Tan Phan, Son Pham

## Abstract

Most widely used tools for finding marker genes in single cell data (SeuratT/NegBinom/Poisson, CellRanger, EdgeR, limmatrend) use a conventional definition of *differentially expressed genes*: genes with different mean expression values. However, in single-cell data, a cell population can be a mixture of many cell types/cell states, hence the mean expression of genes cannot represent the whole population. In addition, these tools assume that gene expression of a population belongs to a specific family of distribution. This assumption is often violated in single-cell data. In this work, we define marker genes of a cell population as genes that can be used to distinguish cells in the population from cells in other populations. Besides log-fold change, we devise a new metric to classify genes into up-regulated, down-regulated, and transitional states. In a benchmark for finding up-regulated and down-regulated genes, our tool outperforms all compared methods, including Seurat, ROTS, scDD, edgeR, MAST, limma, normal t-test, Wilcoxon and Kolmogorov–Smirnov test. Our method is much faster than all compared methods, therefore, enables interactive analysis for large single-cell data sets in BioTuring Browser. Venice algorithm is available within Signac package: https://github.com/bioturing/signac ^1^).

## 1 Introduction

Single-cell RNA-sequencing provides unprecedented insights for cell heterogeneity. At the same time, it yields many new formidable computational challenges. Finding marker genes is among such problems. Most widely used methods for finding marker genes (Seurat-wilcox/t/negbinom/poisson, CellRanger, EdgeR, limmatrend) rely on a conventional definition of *differentially expressed genes* from bulk RNA-seq: *genes with different means of expression values*.

However, single-cell data is very different to bulk RNA-seq data. Generally, a cell population can be a mixture of many cell types, or cell states, hence a single parameter (mean) cannot represent the whole population. For instance, given a population with three sub-populations A, B, C, gene *g* expresses at intermediate, low, and high levels, respectively. This expression patterns often appears in cells at a transitional state (population B) [1]. When this transitional population is compared against the remaining population (cells in A and C), which contain both up-regulated and down-regulated sub-populations, the mean expression value can not be distinguished. Hence, these *transitional genes* can not be revealed by normal *differentially-expressed gene* procedures. In addition, the assumption that gene expression of a population belongs to a specific family of distribution is strongly violated, therefore, the result can be inaccurate.

Some authors had recognized this problem, and incorporated non-parametric tests into their tools, e.g. SigEMD [2], and EMDomics [3], D3E [4] or using a mixture model, e.g. scDD [5]. However, those tools do not have a intuitive statistic like log-fold change to control the quality of the marker genes, and they are also very computational expensive.

In this work, we define marker genes of a cell population as the genes that can be use to distinguish the cells in the population from cells in other populations, and based on that idea, we introduce two simple statistics to quantify the distinctiveness and to classify up/down-regulated or transitional marker genes. This method is also much faster than all compared methods, therefore, enables interactive analysis for large single-cell data sets (up to millions cells).

## 2 Method

We define marker genes of a cell population as genes that can be used to distinguish cells in the population from cells outside of the population. From this idea, we construct the ideal classifier that tries to identify the population of each cell given the expression level of each specific gene. We approximate the classifier’s accuracy and use it as a metric to quantify the "marker quality" of the gene.

### 2.1 Classifier accuracy

Assume that we need to find the marker genes of a specific cell population, denoted as *C*_1_. And we denote the remaining cell population as *C*_2_.

For a gene *g*, let *A*(*g*) be the accuracy of the classifier on identifying the population of the cells, given expression level of gene *g* for each cell. We can split our accuracy into two parts,

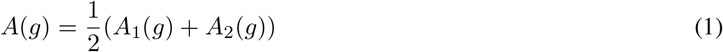

Where *A*_1_(*g*) is the accuracy of identifying the population of cells from *C*_1_, and *A*_2_(*g*) is the accuracy of identifying the population of cells from *C*_2_. In particular,

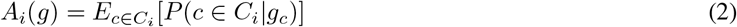

Where *g_c_* is the expression level of gene g in a random cell *c*, and *P* (*c* ∈ *C_i_* | *g_c_*) is the our classifier predicted probability of cell *c* with gene expression *g_c_* belonging to group *C_i_*.

Using the Bayes’ theorem, we have

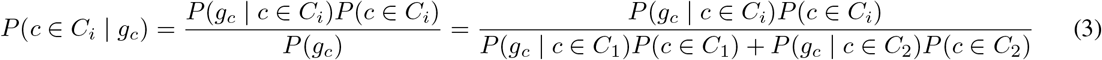

where *i* ∈ {1, 2}.

To avoid the biases toward either population, we simply assign 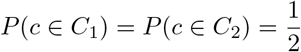. And from (1), (2), and (3), we can deduce the accuracy of our ideal classifier,

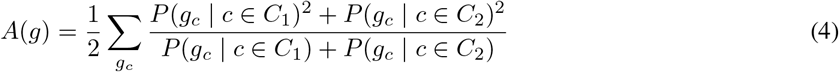

However, *P* (*g_c_* | *c* ∈ *C_i_*) is a latent parameter and the ideal classifier model needs to find its value exactly. As we cannot compute *A*(*g*) directly, we may estimate *A*(*g*) using the estimation of *P* (*g_c_* | *c* ∈ *C_i_*) from our observed data,

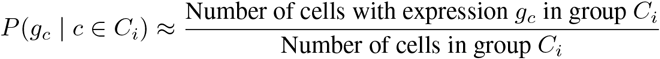

However, using above substitution to estimate *A*(*g*), in many cases, can cause our estimation to have a very high variance. We try to alleviate the problem by grouping the expression level into *k* intervals: *G*_1_, *G*_2_,…, *G_k_*^1^ and denote the new statistic as

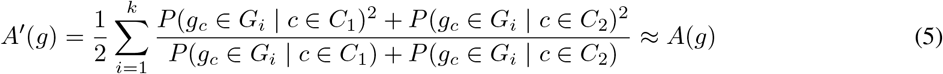

Now, we can substitute the following estimation,

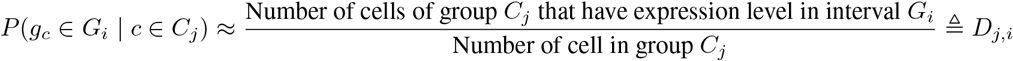

And we have our final estimation of *A*(*g*),

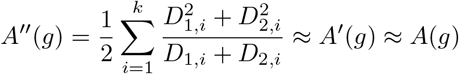

One of the unmentioned problems is that *A*′′(*g*) is a biased estimation of *A*′(*g*). However, as the number of data points in single-cell studies is high, this bias is insignificant. Nonetheless, we provide a correction for the bias, which leads to better score consistency to handle populations with small numbers of cells. We denote the corrected value *A*′′′(*g*), which is defined in the Supplementary.

In our implementation, we rescaled value of *A*′′′(*g*) to 2*A*′′′(*g*) − 1, and called it dissimilarity score. The dissimilarity score can be negative due to bias correction.

### 2.2 Up-down score

Conventionally, we want to classify our differential expressed genes into up-regulated and down-regulated genes. The log-fold change provides a simple metric for the distinction. However, such distinction is only suitable for simple distribution families of gene expression. For genes with more complex distributions, e.g. multi-modal distributions, such distinction cannot always be made.

Instead we propose a different definition of up-regulated and down-regulated genes. We say that a gene is up-regulated in group 1 *iif* for every *p* ∈ (0, 1), the *p*-quantile of the expression in the group 1 is higher than the *p*-quantile of the expression in the group 2 and vise versa for down regulated genes. For genes that do not fit into either cases, the *p*-quantile of the group 1 expression is greater compared to group 2 for *p* ∈ *P*_+_, and smaller for *p* ∈ *P*_−_. For those genes, we assign them the values ranged from −1 to 1 that represent the difference between the measure of *P*_+_ and *P*_−_ and call them *transitional genes*.

Let *Q*_1_, and *Q*_2_ be the quantile function for the expression level of population *C*_1_, and *C*_2_, respectively.

Then, we have the measure of *P*_+_ and *P_−_*,

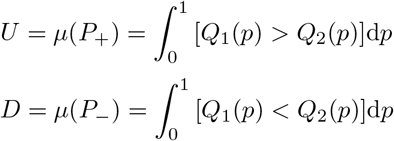

Where 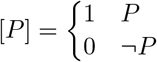.

Finally, we define our up-down score,

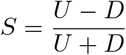

To complete our definition, we need define our quantile function. Given a sorted data set *A* = (*a*_1_, *a*_2_,…, *a_n_*), with *a_i_* ≤ *a_i_*_+1_∀*i* ∈ {1, 2,…, *n* − 1}, we define our quantile function as

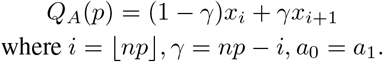

### 2.3 P-value approximation

Our null hypothesis is that the probability that expression level *g_c_* ∈ *G_i_* does not depend on which population that cell *c* comes from.

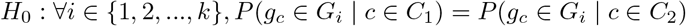

To test the null hypothesis, for each gene *g*, we calculate our statistic *A*′′(*g*) and approximate its p-value using the following theorem (See more details in the Supplement).

#### Theorem P-value theorem

*Under null hypothesis, and fixed intervals G_i_, we have*

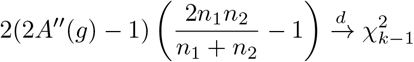

*as the sample sizes n*_1_, *n*_2_ *of both populations tends to infinity*.

*Where* 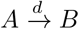 *means the distribution of A converges to the distribution of B*

We also implement a permutation test, which simulates the sampling process from the null sample, in order to estimate the p-value. The permutation test is more accurate in the data with very small cell population. Besides these extreme cases, both methods produce similar results (See the Supplement).

## 3 Results

### Simulated Data

We benchmark Venice precision and performance against 15 other methods using two simulated data sets.

- **Data set 1**: We use scDD, a Bayesian modeling framework, to characterize different expression patterns [6]. In particular, scDD simulates count data from mixtures of negative binomial distribution for two groups of 500 vs 500 cells. The simulated data set contains 10,000 genes, divided into 6 groups: 500 differential expression (DE), 500 differential modality (DM), 500 differential proportion (DP), 500 both differential modality and 500 different component means (DB), 4000 equivalent expression (EE), and 4000 equivalent proportions of cells belonging to each component (EP) (See Figure 1).
- **Data set 2**: To simulate data sets with small number of cells, we sub-sample the above data set to create a new set of 30 samples. Each sample contains two populations, one with 50 cells and the other with 100 cells. Each sample has 1000 differential distribution genes (250 genes for each type of differential distribution) and 3000 equivalent expression genes (1500 EE genes and 1500 EP genes).

scDD classifies differential-distributed genes into four different types (DE, DP, DM, and DB), which are visualized in Figure 1, and non-differential genes into 2 types depending on the modality of the gene expression distribution. EE represents genes with uni-modal distribution and EP represents genes with multi-modal distribution.

**Figure 1:**
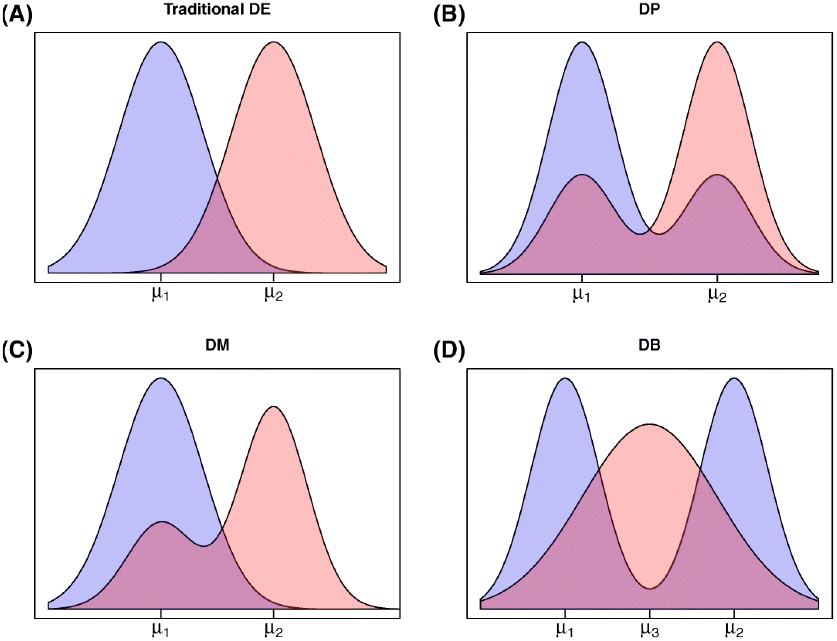
Four types of differential distribution in gene expression. (A) traditional differential expression (DE), (B) differential proportion within each mode (DP), (C) differential modality (DM), and (D) both differential modality and differential expression (DB).

**Figure 2:**
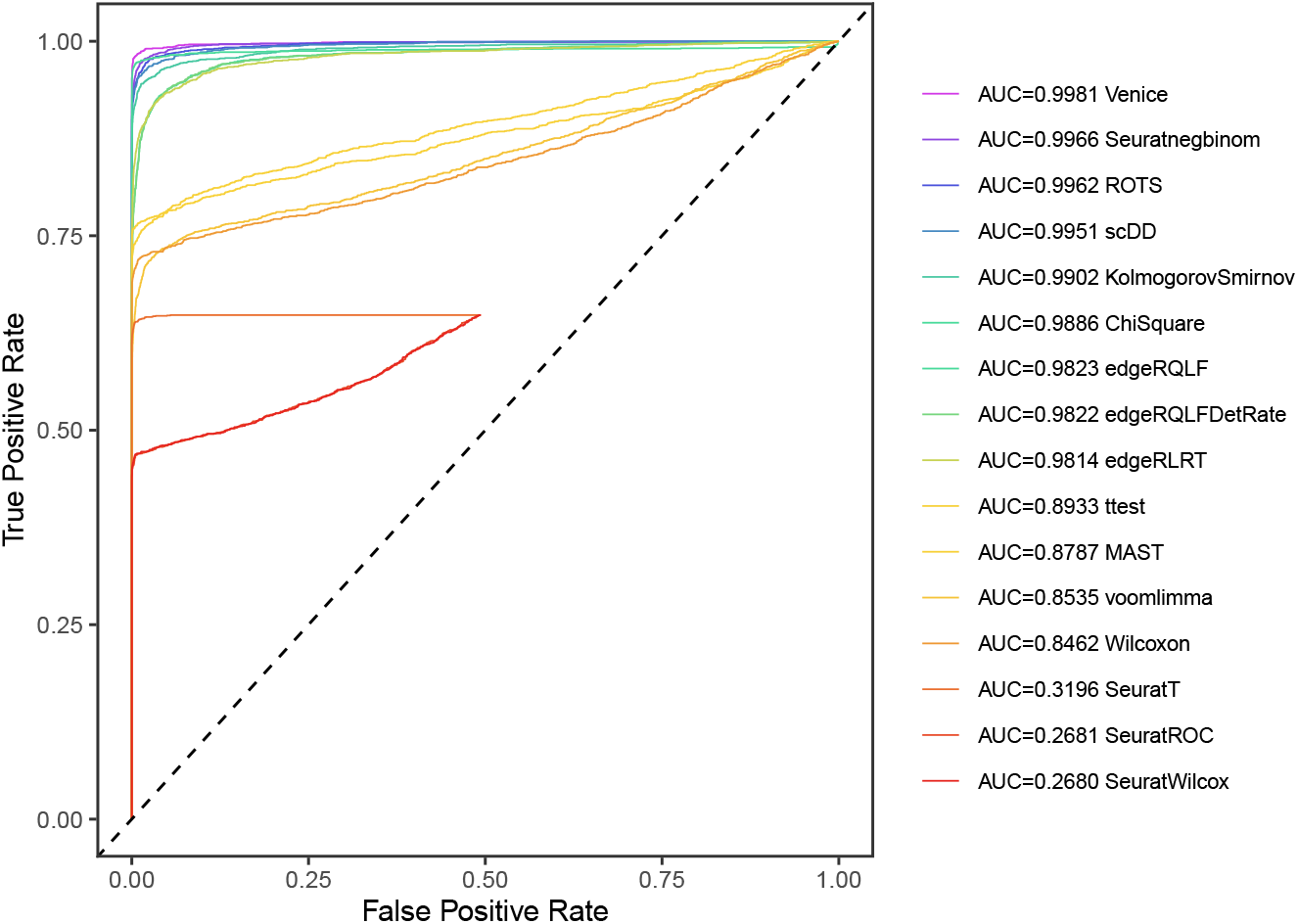
ROC curves of 16 different methods on data set 1. Venice has the best AUC

### 3.1 Data set 1

In the first data set, Venice outperforms all other tools in the benchmark in the AUC score. There is a significant discrepancy between 2 groups of tools. Venice, SeuratNegBinom, ROTS, scDD, KS test, edgeR have AUC scores greater than 0.98, while other tools have AUC scores smaller than 0.9. scDD provided two settings to run. We use the default one, which uses the KS test. The other setting, which uses permutation test for its the Bayes factor, is very computational expensive and doesn’t produce better result in our test. However the KS test of scDD outperforms the standard KS test. SeuratNegBinom also performs really well despite using a simple distribution model. This may stem from the fact that the uni-modal genes are simulated from a negative binomial distribution.

When using the false discovery rate cut-off of 0.05, edgeR methods produce a high number of false positives. Venice and ROTS controls the false discovery rate best as shown in figure 3 and 4. Venice is the tool that detects the most DB genes, which can be the transitional genes.

**Figure 3:**
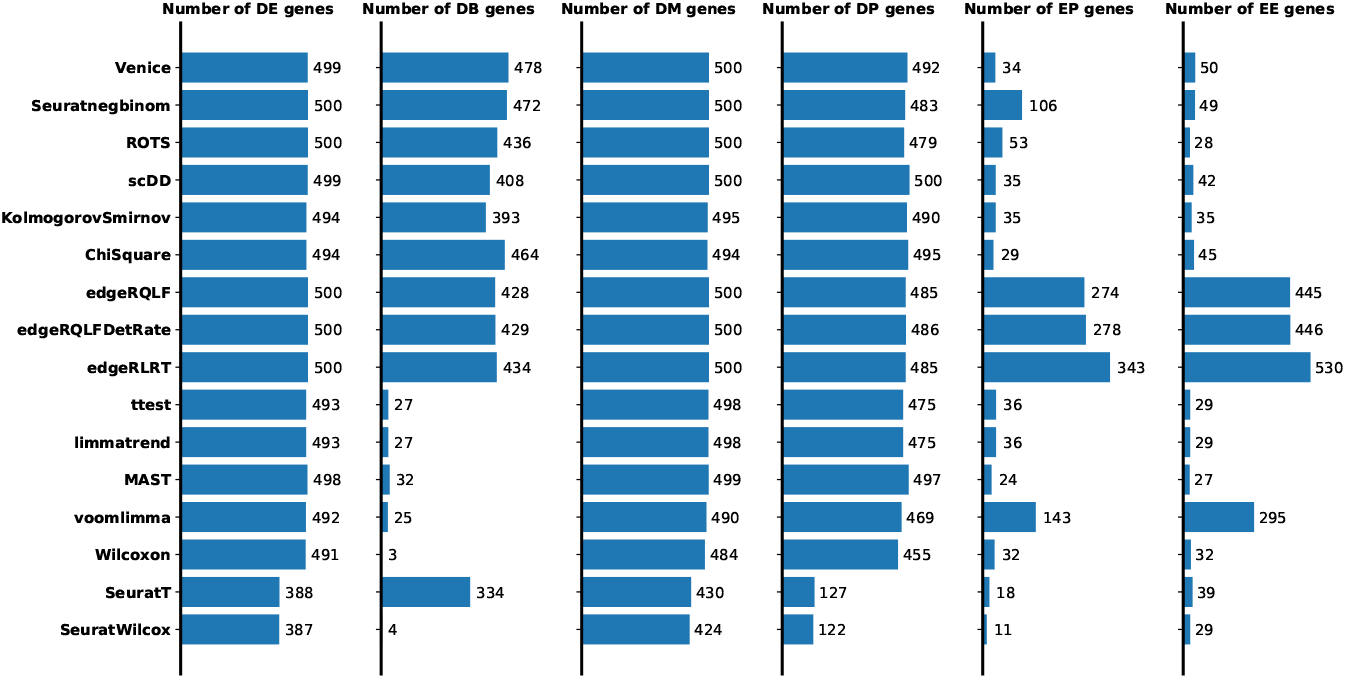
The number of genes in each group that each tool called using a false discovery rate cut-off of 0.05. As explained, DE, DB, DM, DP are true positive; EE, EP are false positive.

**Figure 4:**
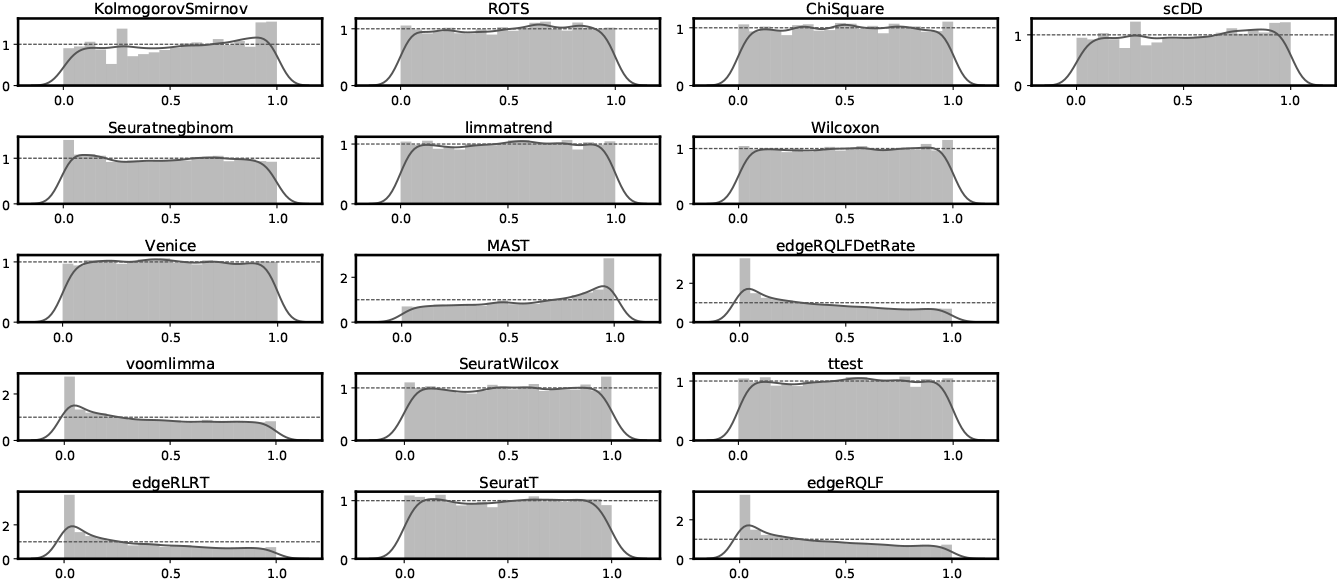
The p-value histogram of non-marker genes of each tool.

**Figure 5:**
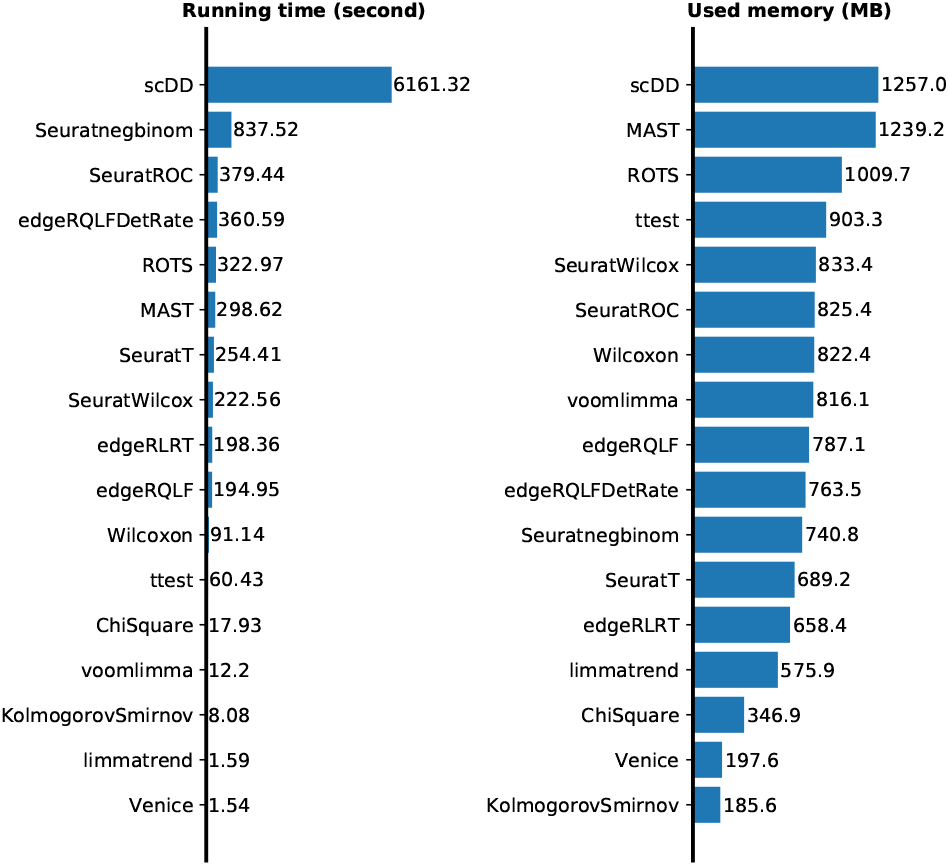
The running time and memory consumption of each tool.

**Figure 6:**
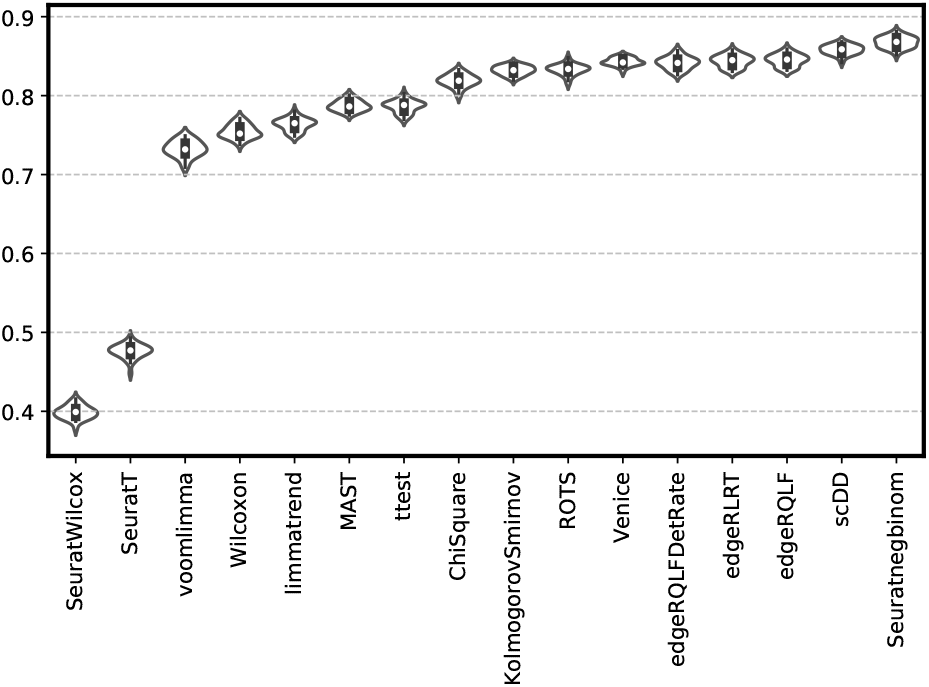
The average AUC of each tool on data set 2.

Amongst the tools with high AUC scores, only Venice and KS test run under 3 minutes on this data set, especially Venice only takes less than 2 seconds. Venice and KS test also use significantly less memory compared to other tools.

### 3.2 Data set 2

In smaller data sets (which are uncommon for single-cell data), our method doesn’t produce the best result. However, the tools with better AUC scores (edgeR, SeuratNegBinom) are assuming the negative binomial distribution. This assumption can improve the power of these tests, however, it causes biases in the p values. Consequentially, it invalidates the false discovery rates. (Figures 7 and 8).

**Figure 7:**
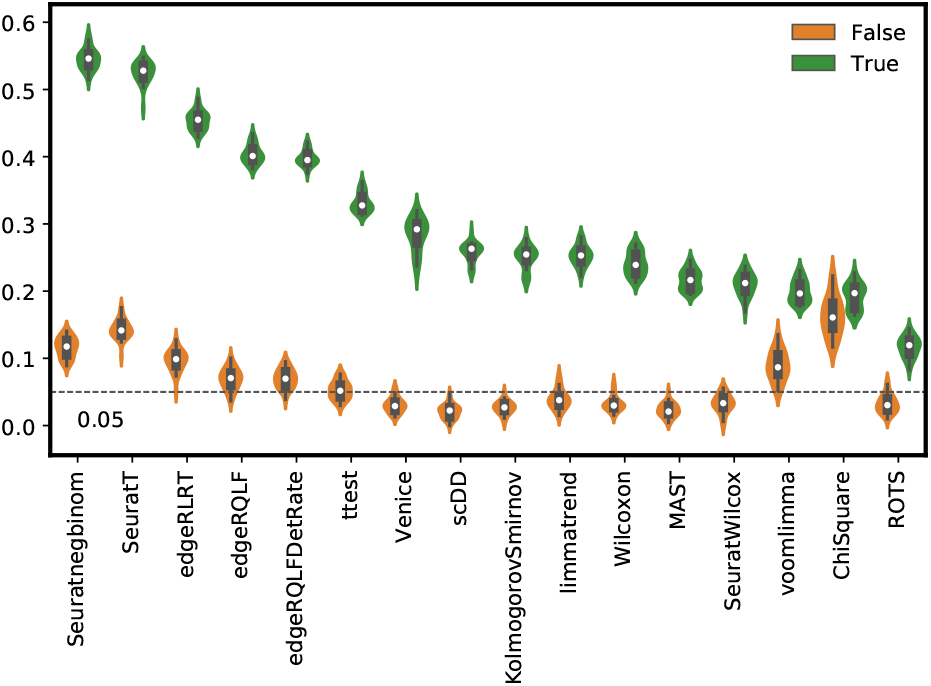
The average false/true positive rate of each tool on data set 2 with the false discovery rate cut-off of 0.05.

**Figure 8.**
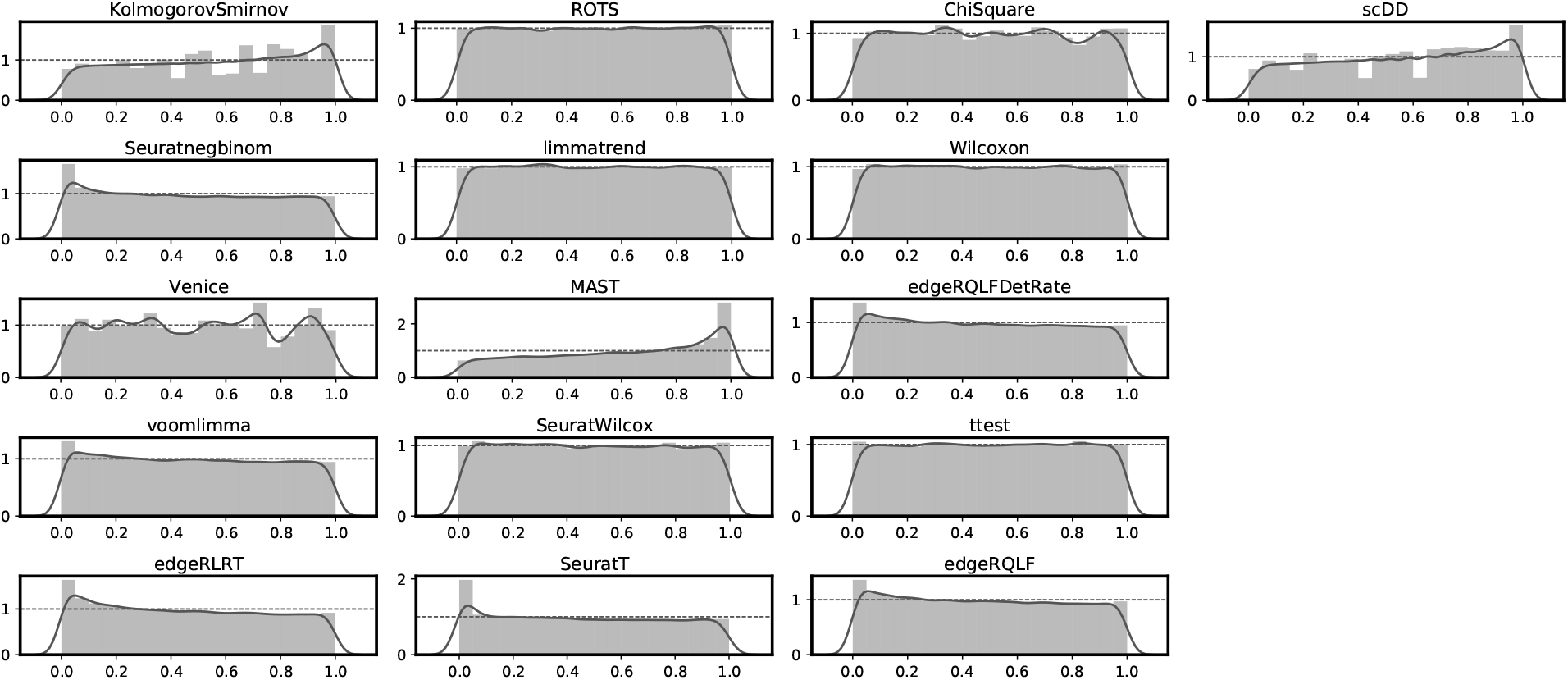

## 4 Supplementary

### 4.1 Grouping

#### 4.1.1 Grouping strategy

First, we want to have about 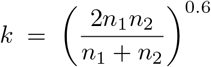 groups. Next, we want each group have a similar amount of cells and we don’t want each group to have too few cells, specifically less than 10 cells. Therefore, we define 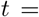 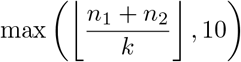 as our expected number of cells in a group.

In our grouping process, first we sort the cells increasingly by gene expression of a gene, we pick out the first *t* cells and put them into a group. However there the next cells may have the same expression level as the *t*^th^ cell. In that case, we also put those cell into that group. We continue our process until don’t have enough cells to form a group. If the number of remaining cells is less than our minimal threshold (10 cells), we merge them into the previous group. Otherwise, we form a new group with our remaining cells.

#### 4.1.2 Effects on the approximation

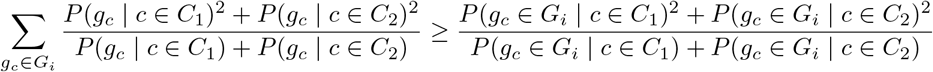

This inequality can be deduced from the Cauchy-Schwarz inequality in Engel’s form 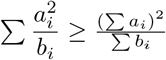

Hence, *A*(*g*) ≥ *A*′(*g*)

Grouping reduces our predicting power, hence causes the accuracy to decrease. However, this is necessary, since there are not always enough data to estimate the expected gene expression for each possible expression level.

### 4.2 p-value

Our main theorem is

#### Theorem 4.1 P-value theorem

*Under null hypothesis, and fixed intervals G_i_, we have*

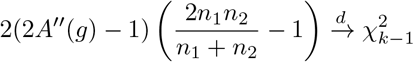

*as the sample sizes n*_1_, *n*_2_ *of both populations tends to infinity.*

Coincidentally, our statistic *A*′′(*g*) is very similar to the Chi-square test statistic after a bit of transformation. Conveniently, to prove this theorem, we can use the same technique that shows Chi-square test statistic follows a chi-square distribution.

First, we needs a few definition.

First we denote *p_i_* = *P* (*g_c_* ∈ *G_i_*)

Under the null hypothesis, it also means that *p_i_* = *P* (*g_c_* ∈ *G_i_* | *c* ∈ *C*_1_) = *P* (*g_c_* ∈ *G_i_* | *c* ∈ *C*_2_)

Next, we denote *D_i_* = (*D_i,_*_1_, *D_i,_*_2_,…, *D_i,k_*) as the observed count vector for population *i*. More specifically, *D_i,j_* is the number of cells of population *i* that have the expression level in interval *G_j_*. We can model *D*_1_ ∼ Multi(*n*_1_, *p*), and *D*_2_ ∼ Multi(*n*_2_, *p*) under the null hypothesis.

And then, we denote the proportion vector 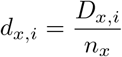 and the average proportion vector 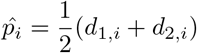

We derived an unbiased estimator *S* of *d*_1_ − *d*_2_ co-variance matrix.

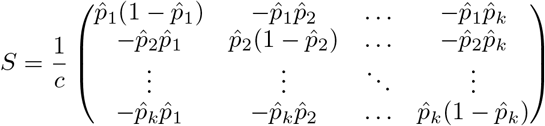

where 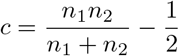

We create *S*\* by removing the last row and column of *S*.

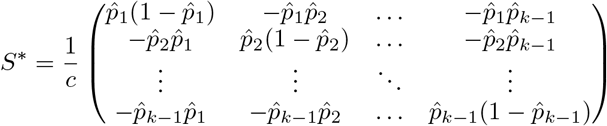

Though *S* is not invertible, as the sum of each column/row is zero, *S*\* is invertible and its inverse can be found using the Sherman–Morrison formula.

Inverse of *S*\*

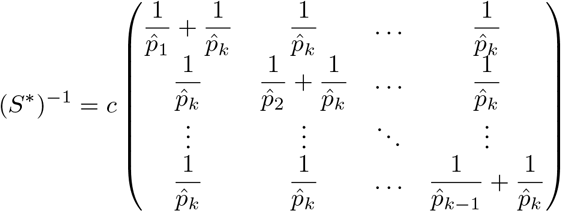

We also denote 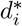 as the vector *d_i_* with the last component removed.

In the following proves, we will ignore the cases when *d*_1,*i*_ = *d*_2,*i*_ = 0 for some *i*, because as *n*, the probability of those cases happening become zero, hence the our theorem below is still hold without explicitly handle those cases.

Before proving the theorem, we need some lemmas. Note that we always assume the null hypothesis and fixed grouping intervals *G_i_*.

#### Lemma 4.2.

*We have* 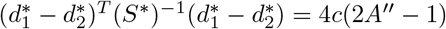

*Proof.* 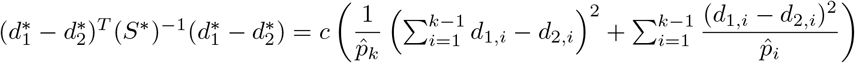

Furthermore, we have that 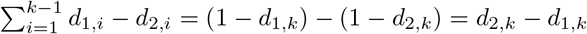, since 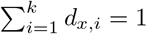

Therefore, we have

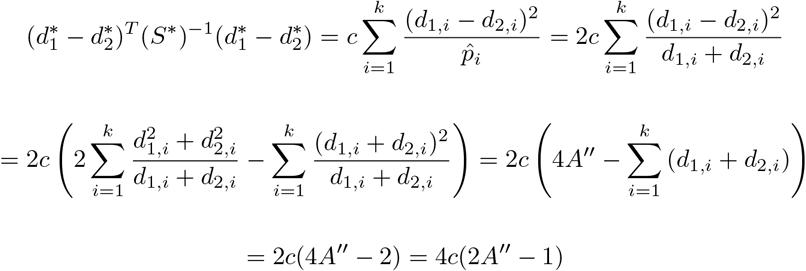

#### Lemma 4.3.

*e have* 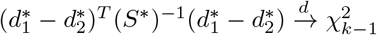 *as n*_1_, *n*_2_ → ∞

*Proof.* First we assume that 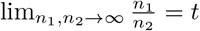 (*t* can be ∞),

Using the central limit and Slutsky’s theorem, we have

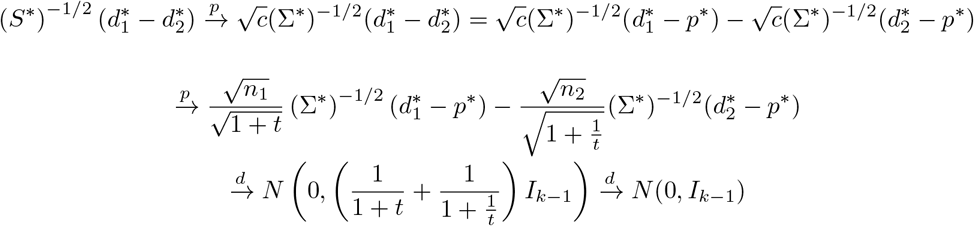

Therefore,

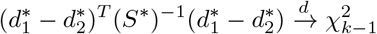

Hence,

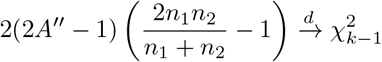

We also need to handle the cases where *t* = 0 and *t* = ∞ to complete this proof.

Here we still have to assume hat 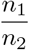 has a limit (maybe infinite). We outline a proof that this assumption is not necessary in the following paragraph

From the Bolzano–Weierstrass theorem, we can show that every sequence of 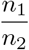 will contain a sub-sequence with a limit (infinite when the sequence is unbounded). Hence, if we only assume (*n*_1_, *n*_2_) → ∞, then for every infinite sub-sequence of (*n*_1_, *n*_2_), we can always find an sub-sub-sequence such that 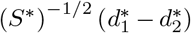 converges to *N* (0, *I*_*k*−1_).We can apply the lemma 4.4 to show that that 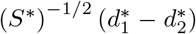 converges to *N* (0, *I_k−_*_1_) without the assumption that 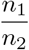 has a limit.

#### Lemma 4.4 Convergence

*Given a sequence* (*a_n_*) *such that every infinite sub-sequence will have a sub-sequence that converges to a common limit L, then the sequence converges to the same limit L*

*Proof.* We will proof this lemma by contradiction.

Assume that the said sequence doesn’t converge to *L*, then by negating the definition of limit, there will exist an *ϵ* > 0, such that there is infinite of *a_n_*, such that |*a_n_* − *L*| ≥ *ϵ*. Then, we construct an infinite sub-sequence from these *a_n_*, denoted (*b_n_*). By the assumption, *b_n_* has a sub-sub-sequence that converges to *L*, hence there exists *b_n_* such that |*b_n_* − *L*| < *ϵ*, which contradicts the construction of *b_n_*.

Here is the proof of theorem 4.1

*Proof.* From the lemma 4.2, we have 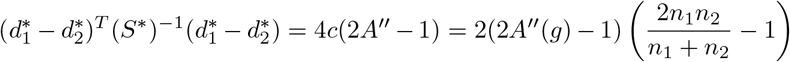

Furthermore, we also have 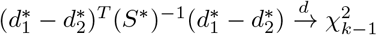 as *n*_1_, *n*_2_ → ∞ from lemma 4.3.

Hence, 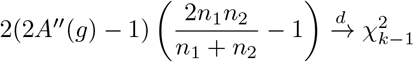

About the fixed grouping assumption, we think it can be loosen up. If the grouping strategy converges to a fixed grouping where every group have a non-zero probability of having a cell when *n*_1_ and *n*_2_ go to infinity, then we think our theorem is still hold.

Though the theorem only provides the asymptotic distribution of the statistic, this distribution seems to approximate the statistic well in common cases.

### 4.3 bias correction

Our estimation of the accuracy is not an unbiased estimator due to grouping the expression level into *G_i_* intervals and the *A*′′ estimation of *A*′. Grouping may cause the accuracy to reduce (*A*′ ≤ *A*), while the *A*′′ cause upward bias (E[*A*′′] ≥ E[*A*′]). The later fact can be shown using the Jensen’s inequality since *A*′ is a convex function of *P* (*g_c_* ∈ *G_i_* | *c* ∈ *C_j_*).

Assume fixed grouping intervals *G_i_*, *D* is a sufficient statistic for the distribution of *A*′. We can assume a estimator *A*′′ of *A*′ is a function of *D_j,i_*. Consequently, E[*A*′′] is a polynomial in *P* (*g_c_* ∈ *G_i_*, | *c* ∈ *C_j_*). On the contrary, *A*′ cannot be represented as a polynomial in *P* (*g_c_* ∈ *G_i_,* | *c* ∈ *C_j_*). Hence there is no unbiased estimator for *A*′.

However, we can do a small correction that is empirically reducing the bias.

First we try to estimate the expected value using the Taylor’s approximation of *A*′′ at *D_j,i_* = *P_j,i_* = *P* (*g_c_ G_i_* | *c* ∈ *C_j_*),

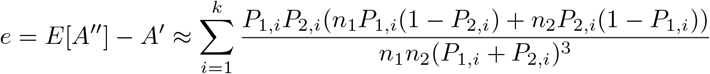

Then, we estimate our bias as,

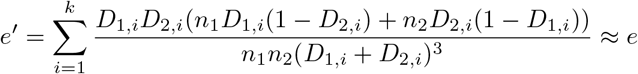

Hence, we have the corrected estimator,

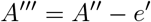

In Venice package, we provide a parameter to turn on and off the correction; it is on by default.

## 5 Acknowledgements

The authors feel very grateful for all the supports from everyone in the BioTuring team, which can range from ideas, suggestions, testings, integration of Venice algorithm to BBrowser, to encouragements or even some pats on the shoulders during some moments when nothing works.

## Footnotes

1 Contact

1 The detail of our grouping strategy is discussed in the Supplementary

## Notes

### Competing Interest Statement

All authors of the paper are supported by BioTuring Inc, a bioinformatics company.

